# Genomic taxonomic assignment of individual root-knot nematodes

**DOI:** 10.1101/2021.03.18.435936

**Authors:** Graham S Sellers, Daniel C Jeffares, Bex Lawson, Tom Prior, David H Lunt

## Abstract

Root-knot nematodes (RKN; genus *Meloidogyne*) are polyphagous plant pathogens of great economic importance to agriculturalists globally. These species are small, diverse, and can be challenging for accurate taxonomic identification. Many of the most important crop pests confound analysis with simple genetic marker loci as they are polyploids of likely hybrid origin. Here we take a low-coverage, long-read genome sequencing approach to characterisation of individual root-knot nematodes. We demonstrate library preparation for Oxford Nanopore Technologies Flongle sequencing of low input DNA from individual juveniles and immature females, multiplexing up to twelve samples per flow cell. Taxonomic identification with Kraken 2 (a *k-mer*-based taxonomic assignment tool) is shown to reliably identify individual nematodes to species level, even within the very closely related *Meloidogyne incognita* group. Our approach forms a robust, low-cost, and scalable method for accurate RKN species diagnostics.

## Introduction

Root-knot nematodes (RKN; genus *Meloidogyne*) are economically important agricultural pests, parasitising all human crops, and causing major losses to world agriculture (Moens, Perry and Starr, 2009). Despite the enormous host range of RKN, perhaps encompassing the diversity of angiosperms (Trudgill and Blok, 2001; Eves-van-den-Akker and Jones, 2018), not all crops are parasitised by all species. Understanding the species identity of individual infections is important in agriculture since it allows understanding of the threat to different crops, better understanding of natural crop resistance, and design and testing of interventions.

Individual RKN juveniles are part of the soil meiofauna and females, attached to or embedded within the root, corms or tubers, are very small (~350 μm), and none of the life stages are straightforward to identify. Species identification may be carried out using microscopy, or with molecular approaches such as isozyme electrophoresis or PCR diagnostics (Blok and Powers, 2009). It can be time consuming to identify RKN, and these diagnostics often do not scale very advantageously in terms of either time or reagent cost. Lastly, although single sample identification is well established, routine testing and understanding of whole nematode communities in the soil is much more challenging. The difficulty is not just because of the greatly increased number of sample assignments required, but also because the diversity of species encountered will increase the taxonomic expertise needed, while simultaneously proving difficult for any DNA barcoding or isozyme approach. Extensive surveys of agricultural environments, their existing nematode communities, and the threat to different planting regimes is therefore largely still an aspiration for pest management.

There are over 90 species of RKN (Karssen, Wesemael and Moens, 2013) and while there is considerable genetic diversity between clades within the genus *Meloidogyne*, there are also species clusters such as the *Meloidogyne incognita* group (MIG) comprising many closely related species including many of the most significant global agricultural pests (Álvarez-Ortega, Brito and Subbotin, 2019).

The MIG, and a number of other species of RKN, are polyploid apomictic asexual parthenogens. They contain divergent A and B subgenomes, which may have either originated from interspecific hybridization or be frozen heterozygosity associated with the loss of sexual reproduction (Dalmasso and Berge, 1983; Abad *et al.*, 2008; Blanc-Mathieu *et al.*, 2017; Szitenberg *et al.*, 2017). Thus, each species contains within every nucleus divergent genome copies that are more distinct from each other than they are from the same subgenome in a different species. This makes it very challenging to use a single loci for species diagnostics since the A subgenome of *M. incognita* is more similar to the A subgenome of *M. javanica* than it is to the B subgenome in the same *M. incognita* cell (see Szitenberg *et al.,* 2017: Figure 3A). Sampling the orthologous loci is therefore essential to reliable identification and understanding of the species, yet this can be very challenging using single locus PCR assays.

Genomic sequencing approaches are able to capture the evolutionary history of species far better than single locus analyses. Genome assembly using short read sequencing technology is very challenging within the MIG, due to our uncertainties about their ploidy, variations in ploidy between strains and the high levels of intragenomic divergence between genome copies. This can pose significant challenges for assemblers, at present. Howerer, long-read high quality data is producing increasingly complete MIG genomes (Koutsovoulos *et al.*, 2020; Susič *et al.*, 2020). Low-coverage long-read Oxford Nanopore Technology (ONT) sequencing can offer a different approach to MIG genomics. For diagnostic analysis, rather than prioritising genome assembly the user could target taxonomic identification, the discovery of novel genetic variability, or the characterisation of virulence-related loci.

Here we develop a method for taxon assignment of RKN using low-coverage, long-read nanopore genome sequencing from single root-knot nematodes. This protocol extracts high quality DNA from juvenile stage 2 (J2), juvenile stage 4 (j4) or immature female individuals, prepares a sequencing library for the Oxford Nanopore Technologies (ONT) Flongle, and allows for multiplexing upto 12 samples per flow cell. From generated sequencing data we use the rapid *k-mer*-based taxonomic assignment tool, Kraken 2 (Wood, Lu and Langmead, 2019). We show that this cost effective, multiplexed metagenomic approach can reliably resolve RKN species including the closely related taxa within the MIG.

## Materials and methods

### Nematode culture

Nematode cultures, established from single egg masses, were grown under quarantine licence on tomato plants (*Lycopersicum solanum*, variety: ‘Moneymaker’) for around 3 months at Fera Science, UK. Single females were excised from the root (with sterile hypodermic needles), rinsed with molecular grade water and frozen in sterile 1.5 ml tubes at −20°C before moving to the University of Hull. J2s were hatched by incubating an isolated egg mass at room temperature in distilled water. Individual j2s were isolated with a dissecting tool, rinsed with molecular grade water and frozen in sterile 1.5 ml tubes at −20°C before moving to the University of Hull.

### DNA extraction

DNA was extracted from individual juvenile stage 2 (j2) or juvenile stage 4 (j4) and immature females excised from root tissue. We employed a modified Mu-DNA: Tissue protocol (Sellers *et al.*, 2018) using a solid phase reversible immobilization (SPRI) magnetic bead capture method, adapted from Rohland and Reich (2012), to isolate high molecular weight DNA (see Supplementary material: Section 2 for detailed protocol).

### Nanopore library construction and sequencing

Library preparation followed a modified version of the ONT Rapid PCR Barcoding Kit (SQK-RPB004). The kit, although at present designed for MinION, was easily converted for use on the Flongle. DNA yield from a single nematode was invariably much lower than the suggested amount for the protocol. However, increasing both the amount of DNA digested and PCR barcoding cycles produced sufficient template for sequencing from all samples (see Supplementary material: Section 3 for detailed protocol).

### Basecalling

Guppy basecaller (version 4.2.2) high accuracy (HAC) GPU was used for basecalling and demultiplexing of MinKNOW (version 20.06.05) fast5 output. Basecalling was performed on the University of Hull’s Viper HPC on a GPU node (2 x 14-core Broadwell E5-2680v4 processors (2.4–3.3 GHz), 128 GB DDR4 RAM, 4x Nvidia Tesla K40m GPU). Reads with a minimum average base quality of 7 and with barcodes at both ends of the read were used for downstream analysis.

### Database construction and cleaning

We used a *Meloidogyne*-specific Kraken2 database as all samples in this study had been taxonomically identified as root-knot nematodes prior to sequencing. This database was considerably smaller (6.5 GB) than a database of all nematode genomes (35.4 GB), and as Kraken 2 loads the database to memory this allowed the analysis to be run locally on a modest computer rather than requiring access to a high memory HPC.

Our *Meloidogyne*-specific Kraken 2 database was constructed from all currently available genomic resources for *Meloidogyne* species from Genbank (listed in Table 1) using scripts provided in the code repository. Despite authors’ efforts to remove contaminant sequences during assembly, some may still remain in the final published genomes. To reduce the effect of these potential contaminants in our analysis, we performed a database cleaning procedure. Genome sequences first underwent a Kraken 2 analysis against a bacterial database (constructed from the Kraken 2 bacteria reference library) with a confidence of 0.1. All unclassified reads from this analysis step were subsequently processed against a custom Kraken 2 database (with a confidence of 0.05) of ‘contaminants’ comprising human (*Homo sapiens*) and a selection of known plant hosts of RKN species; tomato (*Solanum lycopersicum*) and sweet potato (*Ipomoea batatas)*. Genome sequences unclassified at this final step were considered to be free of contaminants and were used for our final database construction.

**Table 1:** Species within the Kraken 2 database. Eleven species of RKN with 21 total genomes are shown In addition to three non-nematode species included to identify contamination.

| Species | Isolate | Accession | Publication |
| --- | --- | --- | --- |
| *Meloidogyne arenaria* | HarA | GCA_003693565.1 | (Szitenberg *et al.*, 2017) |
| *Meloidogyne arenaria* | A2-O Okinawa_01 | GCA_003133805.1 | (Sato *et al.*, 2018) |
| *Meloidogyne arenaria* | Guadeloupe | GCA_900003985.1 | (Blanc-Mathieu *et al.*, 2017) |
| *Meloidogyne chitwoodi* | Race2 | GCA_015183065.1 | unpublished |
| *Meloidogyne chitwoodi* | Race1 | GCA_015183035.1 | unpublished |
| *Meloidogyne chitwoodi* | Roza | GCA_015183025.1 | unpublished |
| *Meloidogyne enterolobii* | Swiss | GCA_903994135.1 | (Koutsovoulos *et al.*, 2020) |
| *Meloidogyne enterolobii* | Sample3 | GCA_903797545.1 | (Koutsovoulos *et al.*, 2020) |
| *Meloidogyne enterolobii* | L30 | GCA_003693675.1 | (Szitenberg *et al.*, 2017) |
| *Meloidogyne floridensis* | SJF1 | GCA_003693605.1 | (Szitenberg *et al.*, 2017) |
| *Meloidogyne floridensis* | JB1 | GCA_000751915.1 | (Lunt *et al.*, 2014) |
| *Meloidogyne graminicola* | VN-18 | GCA_014773135.1 | (Phan *et al.*, 2020) |
| *Meloidogyne graminicola* | IARI | GCA_002778205.1 | (Somvanshi *et al.*, 2018) |
| *Meloidogyne hapla* | VW9 | GCA_000172435.1 | (Opperman *et al.*, 2008) |
| *Meloidogyne incognita* | Kmmt_Gs004 | GCA_014132215.1 | (Asamizu *et al.*, 2020) |
| *Meloidogyne incognita* | W1 | GCA_003693645.1 | (Szitenberg *et al.*, 2017) |
| *Meloidogyne incognita* | Morelos | GCA_900182535.1 | (Blanc-Mathieu *et al.*, 2017) |
| *Meloidogyne incognita* | Morelos | GCA_000180415.1 | (Abad *et al.*, 2008) |
| *Meloidogyne javanica* | VW4 | GCA_003693625.1 | (Szitenberg *et al.*, 2017) |
| *Meloidogyne javanica* | Avignon | GCA_900003945.1 | (Blanc-Mathieu *et al.*, 2017) |
| *Meloidogyne luci* | SS-V13 | GCA_902706615.1 | (Susič *et al.*, 2020) |
| *Homo sapiens* |  | GCA_000001405.28 |  |
| *Tomato* |  | GCA_012431665.1 |  |
| *Sweet potato* |  | GCA_002525835.2 |  |

In addition to the cleaned *Meloidogyne* genomes, we added in a single representative of human, tomato and sweet potato genomes. These extra ‘contaminant’ genomes would allow for evaluation of sample and library preparation from plant host and lab based human contamination.

### Analysis Workflow

We built a reproducible data analysis workflow using the Snakemake workflow manager (Köster and Rahmann, 2012). This workflow uses a conda virtual environment to ensure software compatibility. The basic analysis implements NanoFilt (De Coster *et al.*, 2018) quality control, Kraken 2 (Wood, Lu and Langmead, 2019) taxonomic classification, and presents taxonomic information in textual Kraken 2 outputs and interactive html reports via Recentrifuge (Martí, 2019). The workflow is available at https://github.com/Graham-Sellers/RKN_genomic_taxonomic_assignment.

### Taxonomic assignment method development

Taxonomic assignment methods were developed using a test library (RKN_lib2). This library consisted of 12 *M. incognita* j4 or immature females isolated from root tissue. The samples used ranged from unquantifiable levels of DNA to much higher yields (Supplementary material: Section 5). We considered this to be a relatively diverse library of sample qualities for method development.

#### Quality Control and Taxonomic assignment

Sequencing read quality control was performed using NanoFilt, primarily for head and tail cropping of 50 bp to remove library preparation ligation sites. Additionally, NanoFilt assessed the quality of sequencing reads based on a minimum average read quality score. As our Guppy basecaller parameters were for a minimum average read quality of 7, only quality scores of 7 or above were considered for investigation. Kraken 2 has two parameters that can improve accuracy of read classification: confidence (minimum proportion of *k-mers* required for assignment to a taxonomic level) and minimum base quality score (*k-mers* in the read with bases below this value are considered ambiguous and disregarded for assignment). Optimal values for quality control and Kraken 2 classification parameters were determined by their application across samples in the *M. incognita* test library. These optimised values were used for following investigations.

#### Read counts and taxonomic assignment

We investigated the effect of read number on taxonomic assignment and its effect on the proportion of those reads correctly assigned. Reads were pooled from the 12 *M. incognita* samples. 25 replicates of read count values, at any length, were randomly sampled without replacement from this read pool.

#### Read length and taxonomic assignment

We investigated the influence of sequence read length on taxonomic assignment and its effect on the proportion of those reads correctly assigned. Twenty five replicates of a set read counts for each read length were randomly sampled without replacement from the read pool of *M. incognita*.

#### Display of taxonomy

FIgures 2-5 display the taxonomic assignment and proportion of reads to taxonomic level. These are simplified versions of the Recentrifuge interactive html pages generated by the workflow, and are focused on Meloidogyne and below. Taxonomy follows the NCBI scheme for which some *Meloidogyne* species have a subgeneric taxonomic group “MIG” while others have only genus and species. This NCBI taxonomy is not completely representative of current *Meloidogyne* phylogenetic relationships (Álvarez-Ortega, Brito and Subbotin, 2019), but is still fit for purpose for our species diagnostic method. In order to represent this NCBI taxonomy both our figures, and Recentrifuge when focused on *Meloidogyne*, display 3 concentric rings: the inner one is genus, the outer one is species, and the central ring represents either species or the MIG taxonomic grouping.

### Taxonomic assignment of diverse RKN species

In addition to the M. incognita library, and to increase the diversity of species sequenced, we created a further two libraries of mixed species (RKN_lib3, RKN_lib4 and RKN_lib5) (Supplementary material: Section 5). This resulted in a total of six species being sequenced: j4 or immature female individuals of M. incognita, M. arenaria, M. javanica, M. chitwoodi, M. enerolobii and M. hapla, along with j2 individuals of M. incognita, M. arenaria, M. javanica and M. hapla. Additionally, to test the ability of the method on multi-species samples we created a library where each sample contained two species with a single j2 individual from each (RKN_lib6) (Supplementary material: Section 5). Using the optimal parameters from taxonomic assignment method development, we tested the ability of the method on these species.

### Reproducibility

A detailed protocol for DNA extraction, library preparation and sequencing is available at http://dx.doi.org/10.17504/protocols.io.butanwie and provided openly in the supplementary material of the manuscript repository available at http://dx.doi.org/10.17605/OSF.IO/VA7S2. The repository also includes the Kraken 2 database used in this manuscript, Kraken 2 and Recentrifuge results from our analysis, and a test dataset.

The code repository includes a detailed list of software requirements, suitable for single-step installation with the conda package manager. Bioinformatics analysis steps were performed with the Snakemake workflow manager employing a virtual software environment built from this environment list. Raw sequence data generated in this work has been submitted to the International Nucleotide Databases under BioProject code PRJNA706653. Code repository available at https://github.com/Graham-Sellers/RKN_genomic_taxonomic_assignment.

## Results

### DNA extraction

DNA extraction using the modified Mu-DNA methodology was successful in producing appropriate quality nematode DNA for genomic data. The Mu-DNA extraction technique proved robust and rapid although a standard phenol:chloroform DNA preparation also generated appropriate amounts of DNA in our hands (see Supplementary material: Section 4).

### Library preparation

Genomic sequence data was successfully generated with our modified version of the ONT Rapid PCR Barcoding Kit (SQK-RPB004) using 12 barcodes per flow cell. An example of flow cell read length and count per sample are shown in Figure 1.

**Figure 1.**
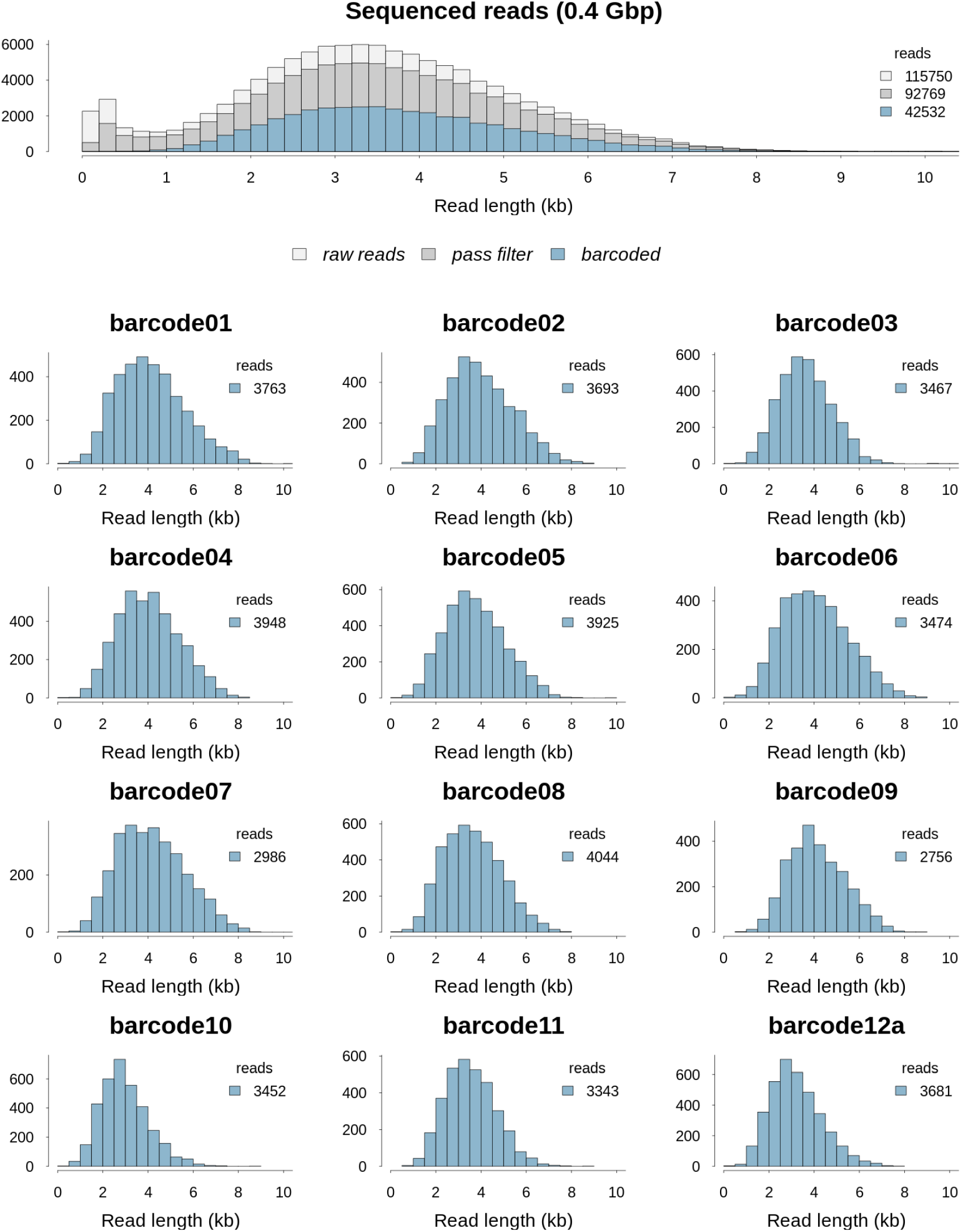
Example of read length distribution from an ONT Flongle flow cell sequencing run. Top panel shows the read length distribution of the sequencing run. Colours indicate the number of raw reads (light grey), reads to have passed filter (grey) and those demultiplexed successfully to barcoded sample (blue). Lower sections show the read length distribution for each barcoded sample. Samples are labeled with raw barcode names.

**Figure 2.**
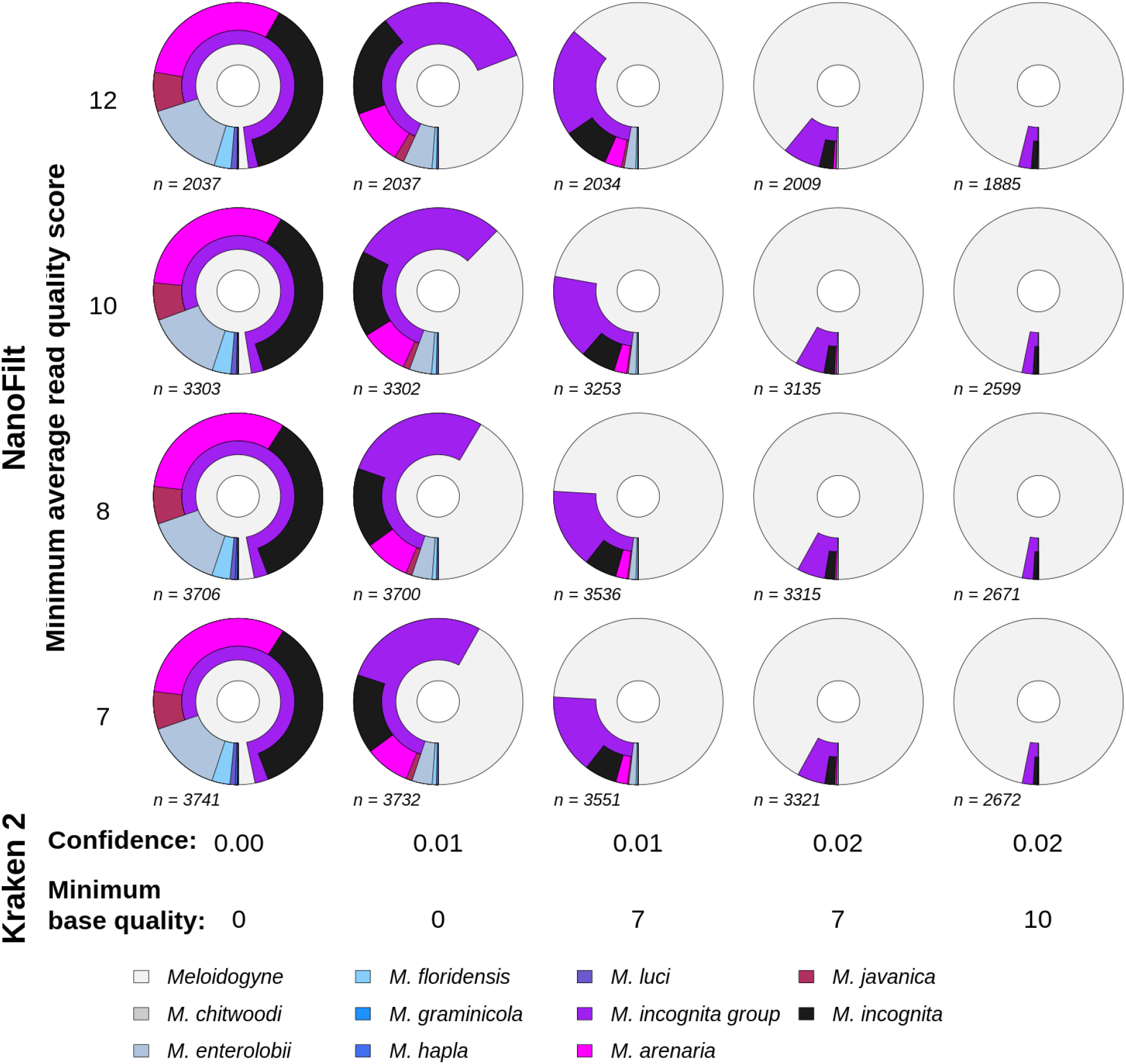
Taxonomic assignment success of *M. incognita* reads varied by quality control and classification parameters. Shown are NanoFilt minimum average read quality scores (y axis) against Kraken 2 confidence and minimum base quality parameter combinations (x axis). Sections show taxonomic assignment of reads (*n*) to the genus *Meloidogyne* and lower taxonomic levels for each configuration of parameters. Each configuration is applied to the same *M. incognita* sample. For each chart the distance from the inner circle represents NCBI taxonomic lineage length; the inner segment is genus level assignment, the second segments are species level assignments (including the MIG species group). MIG species populate the outermost segments, internal to the MIG species group assignments. Segment widths denote proportion reads assigned to that taxonomic level or species.

**Figure 3.**
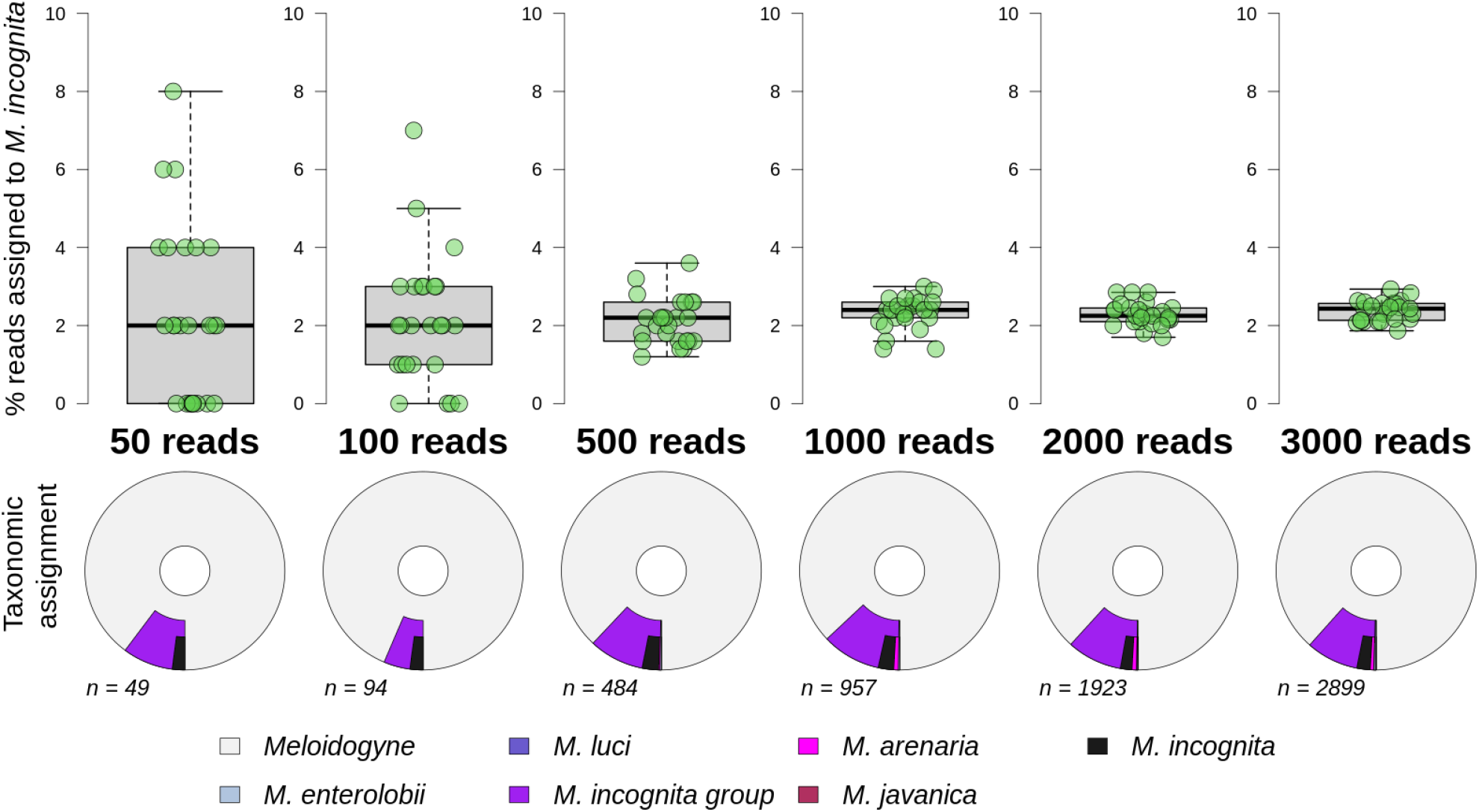
Taxonomic assignment success of *M. incognita* reads varied by read count. Top panel shows the percentage of reads assigned to species level from 50 to 3000 reads. Each treatment has 25 replicates. Lower panel shows a single replicate example of taxonomic assignment of reads (*n*) to the genus *Meloidogyne* and lower taxonomic levels. For each chart the distance from the inner circle represents NCBI taxonomic lineage length; the inner segment is genus level assignment, the second segments are species level assignments (including the MIG species group). MIG species populate the outermost segments, internal to the MIG species group assignments. Segment widths denote proportion reads assigned to that taxonomic level or species.

**Figure 4.**
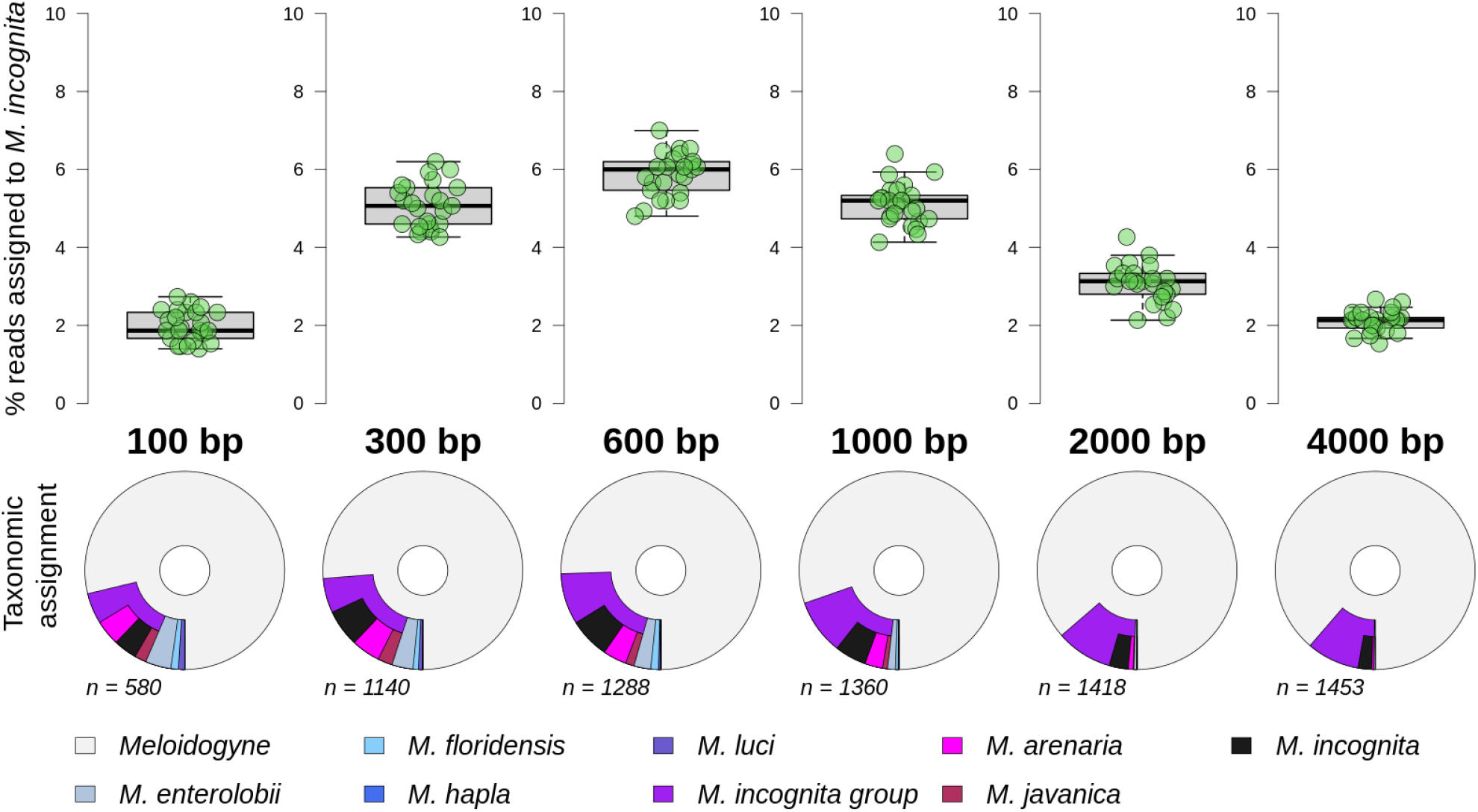
Taxonomic assignment success of 1500 *M. incognita* reads varied by read length. Top panel shows the percentage of reads assigned to species level from 100 to 4000 bp. Each treatment has 25 replicates. Lower panel shows a single replicate example of taxonomic assignment of reads (*n*) to the genus *Meloidogyne* and lower taxonomic levels. For each chart the distance from the inner circle represents NCBI taxonomic lineage length; the inner segment is genus level assignment, the second segments are species level assignments (including the MIG species group). MIG species populate the outermost segments, internal to the MIG species group assignments. Segment widths denote proportion reads assigned to that taxonomic level or species.

**Figure 5.**
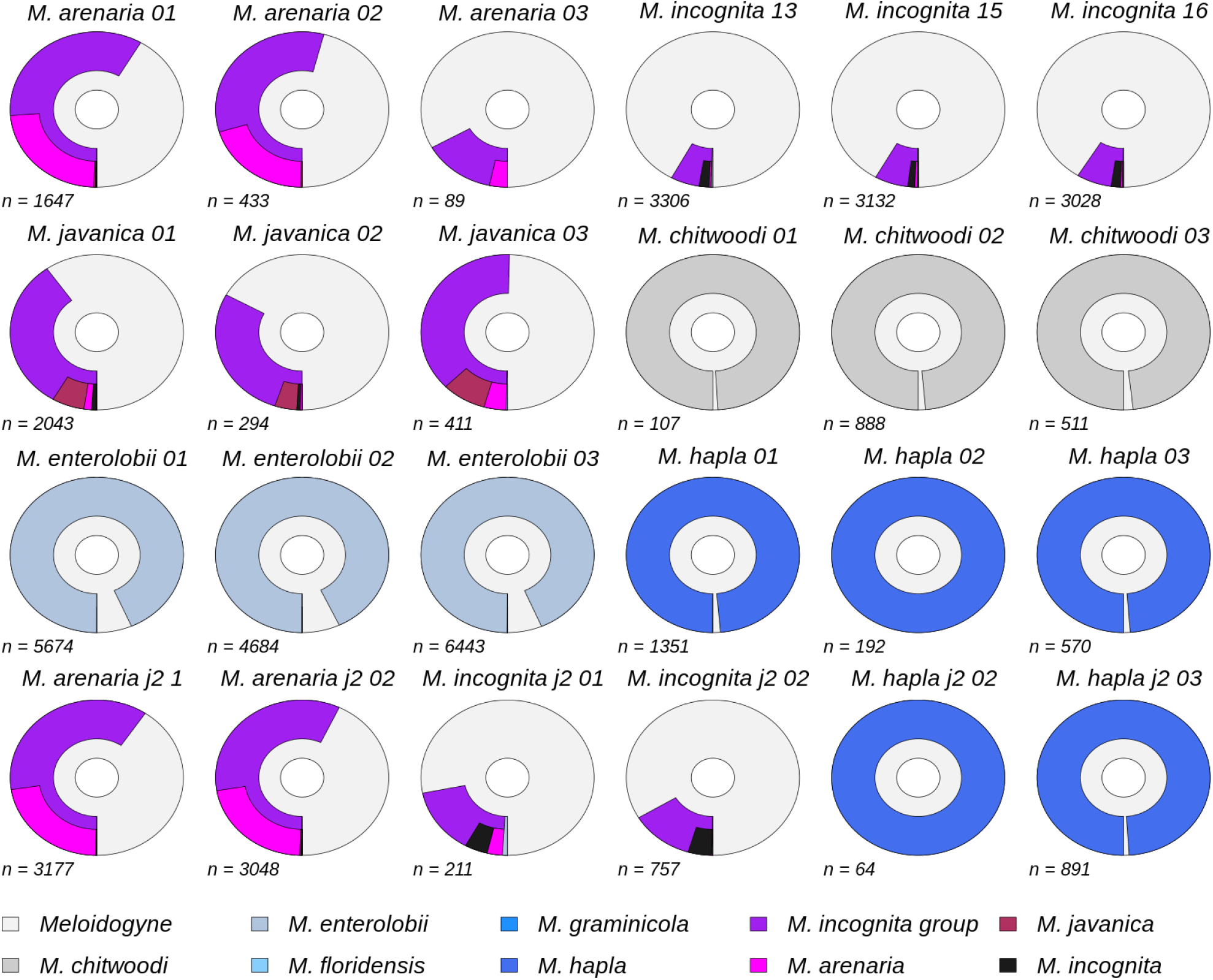
Taxonomic assignment of six *Meloidogyne* species. Top 3 rows are j4 or immature female individuals, bottom row is j2 individuals. Shown are the taxonomic assignment of reads (*n*) to the genus *Meloidogyne* and lower taxonomic levels for individuals of the species. For each chart the distance from the inner circle represents NCBI taxonomic lineage length; the inner segment is genus level assignment, the second segments are species level assignments (including the MIG species group). MIG species populate the outermost segments, internal to the MIG species group assignments. Segment widths denote proportion reads assigned to that taxonomic level or species.

### Database construction and cleaning

The Kraken 2 database we provide was built from currently available RKN genome assemblies with an addition of three non-nematode species (human, tomato and sweet potato) (Table 1). We cleaned the RKN reference genomes of all non-nematode sequences and this removed 380 sequences primarily assigned to bacteria, human and plant. This resulted in a maximum loss of ~1% of sequences from any genome (Supplementary material: Section 6). The database is provided in the manuscript archive but we do not hold it under git version control due to size. Scripts are provided for downloading and formatting databases of RKN genomes, which will be of value to the user since available genomes are increasing continually.

### Taxonomic assignment method development

#### Sequencing output

Read length distributions of our test library (RKN_lib2) are shown in Figure 1, and indicate that overall sequencing produced a mode length of close to ~ 3.5 kb. Successfully demultiplexed sequences per barcode had a mode length of ~ 4 kb (as expected for the library preparation protocol). Low concentration samples (barcodes 9 to 12a) had similar read length distributions compared to higher concentration samples.

#### Quality Control and Taxonomic assignment

Taxonomic identification is largely very robust to the level of sequence read quality control (minimum average read quality score, Figure 2). It is clear that the classification parameters implemented by Kraken 2 have the largest effect. In particular, the most important assignment parameter is the Kraken 2 classification confidence score. However, higher values for read quality control, confidence and minimum base quality considerably reduce the number of reads assigned. To this end we determined that optimal values to reduce noise and remove incorrect assignments were a minimum average read quality score of 7 (i.e. no quality control beyond that of Guppy basecaller) and Kraken 2 confidence score of 0.02 with minimum base quality of 7.

#### Read counts and taxonomic assignment

When multiplexing 12 barcoded samples per Flongle each produced between 3,000 - 4,000 reads per sample (Figure 1). We investigated the effect of smaller read numbers by subsampling 50 to 3000 reads and examining the proportion of reads that were correctly taxonomically assigned. Figure 3 (top panel) shows that the variation in taxonomic assignment success reduces greatly with 500 or more reads, with approximately 2.5% of reads assigned correctly to species rather than a higher level grouping. Assignment was almost exclusively to either the correct species, *M. incognita*, or to the sub-genus level MIG grouping (Figure 3, bottom panel) and this was even true for a randomised 50 read sample.

#### Read length and taxonomic assignment

We investigated the influence of sequence read length on taxonomic assignment. Since our libraries were constructed using the ONT Rapid PCR Barcoding Kit (SQK-RPB004), few reads exceeded 6 Kb. We examined read lengths from 100 bp up to 4 kb, with each treatment having 1500 reads. Figure 4 (top panel) shows that different lengths vary between approximate 2-6% of reads assigned to the correct species (*M. incognita*), with most for the 600 bp reads. However, as shown in Figure 4 (bottom panel) the most frequent species level taxonomic assignment was to the correct species. In the case of the *M. incognita* samples, the major effect of increasing read lengths was to remove the noise of reads identified to any source other than *M. incognita* or the *M. incognita* group. Longer reads have more *k-mers* and has the effect of pulling back noise and allows for an increase in reads classified. Reads longer than 1000 bp were decided as optimal for taxonomic assignment.

### Taxonomic assignment of diverse RKN species

Six species of RKN were sequenced over multiple libraries; *M. incognita*, *M. arenaria*, *M. javanica*, *M. chitwoodi*, *M. enerolobii* and *M. hapla*. Within these samples were juvenile stage 2 (j2) individuals. Reads were not always evenly distributed across samples in a library. Although different nematode samples produced different read counts and qualities, in all cases, using optimal parameters (reads > 1000 bp, Kraken 2 confidence score of 0.02 and minimum base quality of 7), the most frequent assignment to the species level was the correct taxonomy (Figure 5). Species within the MIG had many reads assigned to genus level (*Meloidogyne*) and fewer to species level. In contrast, species outside the MIG (*M. chitwoodi*, *M. enerolobii* and *M. hapla*) invariably had the vast majority of reads assigned to species. Although J2 samples had no quantifiable input DNA they sequenced successfully and had taxonomic assignments similar to their j4 counterparts.

Taxonomic assignment of multi-species samples, consisting of j2 individuals from two species (RKN_lib6), had mixed success. Samples containing two MIG species were difficult to disentangle, whereas those with an individual external of the MIG, e.g. *M. hapla*, had clear assignments to species level for both species (see RKN_lib6 results in the manuscript repository at http://dx.doi.org/10.17605/OSF.IO/VA7S2).

Under optimal circumstances for 12 multiplexed samples, with 3000 reads (assigned to *Meloidogyne* and lower) with an average read length of 4 Kb, we would expect to obtain 12 Mb per sample. This is low coverage, approximately 0.1x MIG haploid genome, and will be lower for samples that have low read counts assigned and considering PCR duplication. Despite this, we have achieved very accurate taxonomic classification of samples with low read counts (Figure 5), and therefore low genome coverage.

## Discussion

Here we demonstrated the effectiveness of a genomic approach as a method for the taxonomic assignment of a single RKN individual. Efficient DNA extraction in combination with optimised library preparation generated long-read sequencing data from ONT Flongle devices. These long-read sequences contained sufficient resolution to successfully diagnose a sample to species, even those within the closely related MIG. Genomic approaches to study RKN typically include culturing large amounts of individuals. Here, in contrast, we show that a single j2 RKN can yield sufficient DNA for reliable identification through our method.

### Genomic approaches

Genetic characterisation of RKN has often been carried out by the PCR amplification of single or a small number of loci. These approaches can be informative but have the disadvantage that they cannot sample and represent the two distinct genomes present in some RKN species, especially the MIG. The presence of both A and B subgenomes, shared across species, requires exceptional vigilance that homologous copies of the genome are being compared, else erroneous results could be obtained.

Genomics samples across the genome, including the A and B subgenomes of MIG. Since *k-mers* are short and exceptionally numerous they will not suffer from problems of A/B subgenome orthology but rather represent a database of oligonucleotide variants. So, unlike PCR assays, both subgenomes are sampled and analysed with no requirement for complex subgenome assignment.

We set out to develop a low-coverage sequencing approach for taxonomic ID from whole genomic DNA. The Flongle protocol that we developed returned a low genomic coverage, though this is unlikely to be an important metric when genome assembly is not involved. Instead an approach which effectively samples from the complex polyploid genome of MIG is likely more important to taxonomic ID. Our method, taking this approach, can be seen to accurately identify samples to species (Figure 5). Read counts assigned to *Meloidogyne* varied greatly between samples, but even in low read count samples the represented genome data therein had sufficient resolution for taxonomic classification. Our approach potential for samples containing more than one species. The results indicated that this might be a powerful approach for separating more genetically distinct taxa in mixed samples. It also showed promise for more closely related species (e.g. within the MIG), however its success here will almost certainly improve with the increasing completeness of the reference database.

### Scalability, cost effectiveness and time eficiency

The ONT Flongle approach developed here allows small scale analysis of up to 12 samples at a time. This is a scale that may fit well with many small laboratories. The protocol may however be scaled up considerably across multiple MinION or GridION devices with Flongle adaptors. While the ONT Flongle can generate up to 2 Gb of data, due to difficulties accessing the laboratory during the COVID-19 pandemic many of the flow cells used in our analysis were considerably past their expiration date and only produced ~ 0.5 Gb data. Despite this we were still able to produce sufficient reads for our analysis process. It is likely that future experiments or different users would yield considerably more data per barcoded sample.

The Flongle genomic approach has costs similar to other types of genetic analysis. Currently a single flow cell, reagents and consumables needed for sequencing 12 samples equates to less than 20 GBP per sample (excluding VAT and start time) (Supplementary material: Section 7). Our method can produce results within as little as 36 hours start to finish, with less than 4 hours expert hands-on time and a 24 hour sequencing period (Supplementary material: Section 7).

### Taxonomic assignment of closely-related MIG species

When looking at *M. incognita*, one of the most challenging species to identify via genetic methods, many isolates are assigned not just to genus or MIG but to species, regardless of low read counts (see Figure 5). These assignments reveal the strength of that result with the percentage of reads being assigned to species and taxonomic levels.

Identification of more than one species in a single nematode preparation does not indicate hybridization or mixing of samples but rather is what would be expected with the genetic similarity of MIG species in addition to sequencing error and an imperfectly resolving database. Most reads are assigned to genus level, indicating the close relationship between species, where *k-mers* do not assign reads to species unambiguously. Despite this, we found most isolates to have a strong assignment to the correct species with a minimal percentage of reads assigned incorrectly.

Our analyses of quality control, read number, and read length indicate that taxonomic assignment of *M. incognita* is largely robust to variation in these parameters and may be accomplished on samples that have not sequenced well. We do identify however that approximately 1000 reads ≥ 1000 bp is a powerful dataset with which to assign taxonomy in the MIG.

### Improving genomic databases improves analysis

Our study tested the identification of RKN species where the species was represented in the database. It is clear however that this will not always be the case for samples, and improving reference databases will be very advantageous to future diagnostics. We suggest expanding the public sequence databases for root-knot nematodes as a priority for our research community as a whole.

Creating genomic resources currently requires sample culturing, whole genome sequencing and assembly. In this respect, it will be interesting to see how well light-coverage, long-read sequencing from single nematodes can meaningfully supplement the genomic database. For example, if single, well-identified individuals from a species are sequenced on an ONT Flongle, would this be sufficient to include in the reference database with a minimal genome assembly method?

### Functional genomics and nematode virulence

A goal of root-knot nematode genomics has been to better characterise the genes present, better understand the physiology and biochemistry of the nematode life cycle, and the likely contribution of key genes to virulence (Abad *et al.*, 2008; Opperman *et al.*, 2008; Bird *et al.*, 2009). Significant progress in this direction is being made (Bird *et al.*, 2015; Castagnone‐Sereno *et al.*, 2019; Cox *et al.*, 2019; Grynberg *et al.*, 2020; Ste-Croix *et al.*, 2021) and this knowledge may soon more widely inform our understanding of the virulence of field isolates of RKN. Although the experimental design implemented here kept the sequence coverage per isolate very low, it would be straightforward to increase this by reducing the number of samples per Flongle flow cell, or upscaling to a MinION flow cell. It is likely that such assays will, in future, informatively report the variation of virulence target loci from single individuals, or even population samples.

## Conclusions

We show that a low-coverage, long-read genomics approach can be used to reliably identify root-knot nematodes. Species identification was robust to much of the variation in read length and read number that we encountered. Our genomic approach is put forward as an accurate, low-cost, and scalable approach to species diagnostics. In addition, we have highlighted the importance of extending the number of agriculturally relevant species for which genomic data is in the public databases. Finally, we allude to how the increasing understanding of the genetic basis of nematode virulence, in combination with rapid individual-based genomics, could deliver more informed assessments of agricultural risk.

## Supporting information

Sequences have been submitted to the International nucleotide sequence databases with BioProject accession number PRJNA706653. All code for data analysis is available under an MIT licence at https://github.com/Graham-Sellers/RKN_genomic_taxonomic_assignment. The Kraken database, results, test dataset, and supplementary material are part of the manuscript reproducible archive, and can be found at http://dx.doi.org/10.17605/OSF.IO/VA7S2. The DNA extraction, library preparation, and sequencing protocol is available under a Creative Commons Attribution licence at http://dx.doi.org/10.17504/protocols.io.butanwie.

## Acknowledgements

This work was funded by a ‘THYME - mobilising the bioeconomy’ award, held at the University of Hull’s Energy and Environment Institute, UK. Thanks go in particular to Jenny Spear and Dan Parsons for support in the development and implementation of this project. We would also like to thank Dr. Robert K. Donnelly for laboratory assistance throughout the project. We acknowledge the Viper High Performance Computing facility of the University of Hull and its support team in achieving this work.

